# Combination of Volasertib and Rapamycin Inhibits the Regrowth of TSC2-Deficient Tumors

**DOI:** 10.1101/2022.11.02.512640

**Authors:** Matthildi Valianou, Erik Y. Zhang, Daniel L. Johnson, Rhett Meyers, Maciej M. Markiewski, Magdalena Karbowniczek, Lawrence M. Pfeffer, Jane J. Yu, John J. Bissler, Aristotelis Astrinidis

## Abstract

Mutations in *TSC1* and *TSC2* lead to hyperactivation of mTORC1 and cause Tuberous Sclerosis Complex (TSC) and pulmonary Lymphangioleiomyomatosis (LAM). Rapamycin and rapalogs are potent inhibitors of mTORC1 activity and are approved for the treatment of TSC and LAM. Nevertheless, rapalogs do not cause tumor cell death, and cessation of therapy leads to tumor regrowth. Polo-like kinase 1 (PLK1) interacts with and phosphorylates TSC1, and PLK1 inhibition induces apoptosis and attenuates autophagy in TSC1/TSC2-deficient cells. Here we report that the PLK1 inhibitor volasertib decreases the viability and survival and induces apoptosis in 621-101 cells, a TSC-deficient renal angiomyolipoma cell line from a LAM patient. Combined with rapamycin, volasertib further decreased the survival of 621-101 cells. *In vivo*, volasertib reduced short-term mouse lung colonization by TSC2-deficient cells and decreased the growth of TSC2-deficient subcutaneous tumors. Mice treated with a combination of volasertib and rapamycin had slower tumor relapse after discontinuation of treatment, compared to rapamycin only. Approximately 35 days after discontinuation of treatment, we observed persistent apoptotic markers and gene expression changes for type I interferon signaling in the regrowth tumors from combination-treated mice, compared to rapamycin only. Notably, combination treatment potently inhibited the growth of subcutaneous tumors derived from the rapamycin-refractory cell line ELT3-245. Taken together, our current work demonstrates that for the management of TSC and LAM disease combination of mTORC1 and PLK1 inhibitors in some cases may be advantageous over currently used rapalog monotherapies.

## INTRODUCTION

The PI3K/mTOR signaling pathway is a critical component of cell homeostasis, integrating extracellular and intracellular cues and pairing them with anabolic processes, such as protein synthesis and biosynthesis of ribosomes and nucleotide precursors, while inhibiting catabolic processes like autophagy. Mutations in key tumor suppressors and oncogenes of this pathway, including *PTEN, PI3K, AKT, TSC1* and *TSC2*, lead to the aberrant regulation of PI3K/mTOR and various cancers (1). The mechanistic target of rapamycin (mTOR) is a serine/threonine kinase that forms two protein complexes, mTORC1 and mTORC2, with distinct components, substrates, and functions. Loss-of-function mutations in *TSC1* and *TSC2* lead to the formation of hamartomatous growths in Tuberous Sclerosis Complex (TSC) and pulmonary Lymphangioleiomyomatosis (LAM) (2-4). Hamartin and tuberin, encoded by *TSC1* and *TSC2*, respectively, form a multimeric protein complex which is regulated by kinases responsive to growth factors, such as AKT and ERK1/2, intracellular energy levels and reactive oxygen species, like AMPK, and by kinases that are regulated during the cell cycle, such as CDK1/cyclin B, PLK1 and PLK2 [(5-8) and reviewed in (9)].

Although rapamycin and its analogues (collectively rapalogs) are approved for the treatment of TSC and LAM lesions, they do not eliminate the mutant cells, discontinuation of treatment leads to tumor regrowth in most cases, and some patients do not adequately respond to treatment (10-12). The main limitations of rapalogs is their cytostatic - rather than cytotoxic - activity, and the activation of pro-survival signaling, mainly the PI3K/AKT (13-15) and autophagy (16), during therapy. Thus, the identification of new druggable targets that interplay with the PI3K/mTOR signaling pathway may result in new and improved therapeutic regimens. Ideally, these therapies should aim to eradicate tumors in TSC and LAM patients.

Polo-like kinases is a family of serine/threonine kinases comprised of five members in vertebrates with diverse functions in the regulation of cell division [reviewed in (17)]. The most extensively studied family member is polo-like kinase 1 (PLK1) which regulates centrosome maturation, mitotic entry, spindle assembly, anaphase, and cytokinesis. Ablation or prolonged inhibition of PLK1 induces formation of monopolar mitotic spindles and prometaphase arrest; these unresolved mitoses lead to mitotic catastrophe and apoptosis (18, 19). Increased levels and activity of PLK1 have been found in various cancers, and several inhibitors are currently being evaluated in clinical trials for oncology. Previously we found that at least two members of the polo-like kinase family, PLK1 and PLK2, interact with the TSC1/TSC2 complex (6, 7), and that the PLK1 inhibitor BI-2536 reduces the viability and survival and induces apoptosis in TSC1- and TSC2-deficient cells, compared to controls (20). In the present study we explore if volasertib (BI-6727), a clinically relevant PLK1 inhibitor (19), can overcome the hurdles associated with rapalog therapy in TSC and LAM.

## RESULTS

### Volasertib decreases the viability and clonogenic survival and induces apoptosis in TSC2-deficient patient angiomyolipoma cells

We previously showed that the PLK1 inhibitor BI-2536 decreases the clonogenic survival of HeLa cells after silencing of TSC1 or TSC2 and of *Tsc1*^-/-^ mouse embryonic fibroblasts (MEFs), compared to TSC1- or TSC2-expressing control cells (20). Additionally, BI-2536 induces apoptosis preferentially in vector-transduced TSC2-deficient ELT3 and *Tsc1*^-/-^ MEFs, compared to those with ectopic expression of TSC2 or TSC1, respectively. To test the efficacy of PLK1 inhibition in a clinically relevant setting we used the human TSC2-deficient cell line 621-101 and volasertib (BI-6727), a PLK1 inhibitor in phase II/III clinical trials (19). 621-101 cells are derived from a renal angiomyolipoma of a sporadic LAM patient (21). Volasertib and BI-2536 decreased the viability of 621-101 cells after a 48 h treatment, with median-growth inhibition concentration (GI_50_) of 2.48×10^−8^ M (24.8 nM) and 7.78×10^−9^ M (7.78 nM), respectively (**Figure 1A, Supplemental Figure S1**, and **Supplemental Table 1**). These data are in close agreement with the average GI_50_ for the two compounds tested against the NCI-60 cancer cell line panel (NCI Developmental Therapeutics Program). Next, we tested the effect of volasertib and BI-2536 on the clonogenic survival of 621-101 cells after 72 h of treatment. Both drugs significantly reduced the absolute number of colonies in all three concentrations tested (3, 10 and 30 nM), compared to DMSO-treated cells (*P* < 0.01 for all treatments vs. DMSO, **Figure 1B**, and **Supplemental Table 2**). In agreement with our viability data and those from the NCI showing a lower GI_50_ for BI-2536, compared to volasertib, 10 and 30 nM BI-2536 caused a significantly stronger inhibition of 621-101 clonogenic survival, compared to the same concentration of volasertib (**Supplemental Table 2**).

**Figure 1.**
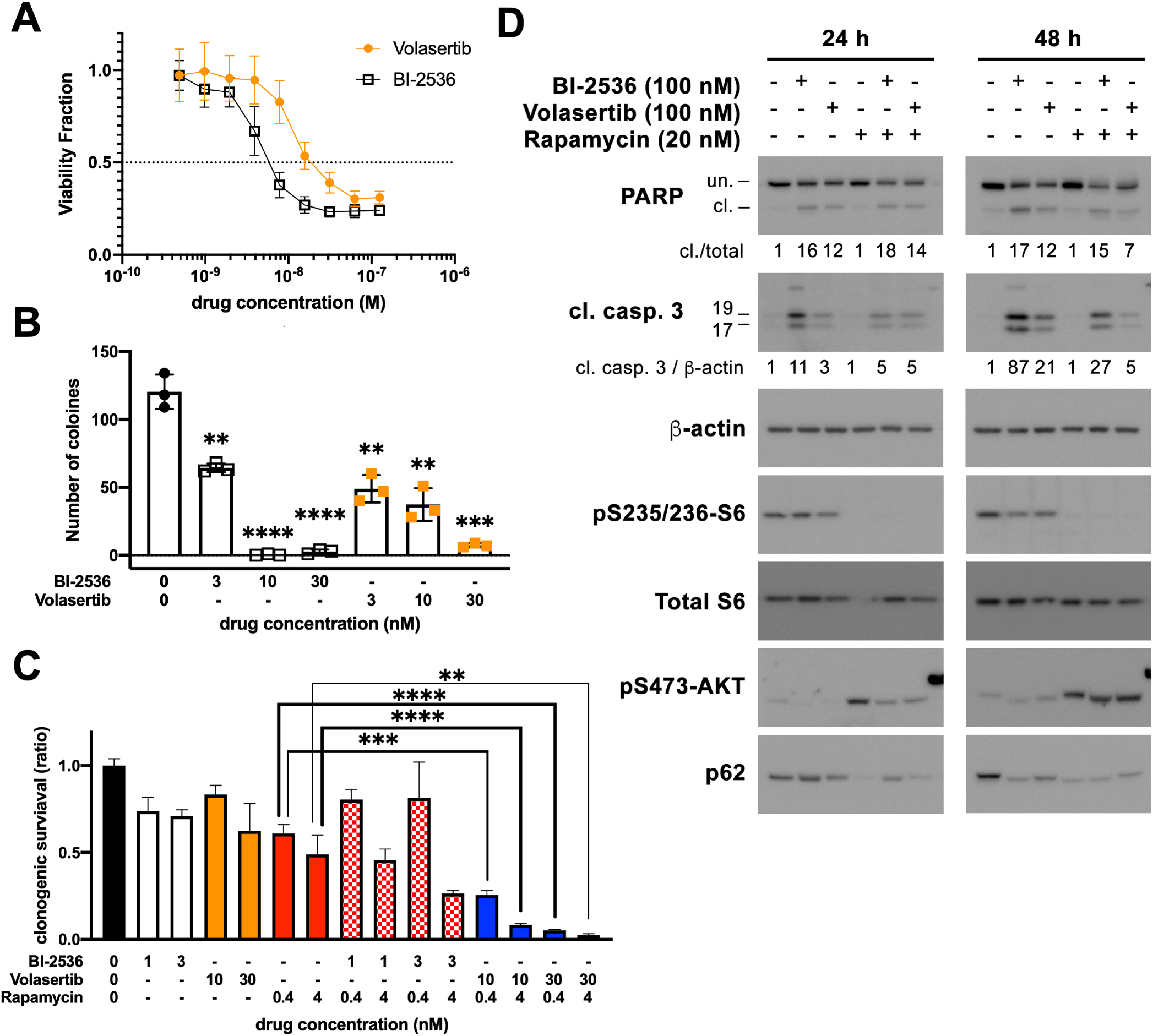
PLK1 inhibitors decrease the viability and survival and induce apoptosis in 621-101 cells. **(A)** Viability of 621-101 cells that were treated with compounds for 48 h. Error bars are SD. **(B)** Clonogenic survival of 621-101 cells after 72 h treatment with compounds or DMSO (vehicle control). Asterisks indicate statistical significance compared to vehicle control treated cells (unpaired T-test). Error bars are SD. **(C)** Clonogenic survival of 621-101 cells after 72 h treatment with compounds, combinations, or DMSO (vehicle control). Clonogenic survival shown in *y* axis is relative to DMSO-treated cells. Asterisks indicate statistical significance between comparison groups (unpaired T-test). Error bars are SEM. **(D)** Immunoblotting of lysates from 621-101 cells that were treated for 24 h and 48 h with compounds, combinations, or DMSO (first lane of each blot). For PARP, the 115 kDa (uncleaved, unc.) and 85 kDa (cleaved, cl.) fragments are shown. Numbers under the PARP immunoblots show the ratio of the cleaved (85 kDa) PARP fragment intensity to the sum of intensities for both PARP fragments (115 kDa + 85 kDa), normalized to the control lysates in the first lane of each immunoblot. For cleaved caspase 3 (cl. casp. 3), both the 19 kDa and 17 kDa fragments are shown. Numbers under the cleaved caspase 3 immunoblots show the ratio of the sum of intensities for both cleaved caspase 3 fragments (19 kDa and 17 kDa) to the intensity of the β-actin band, normalized to control lysates in the first lane of each immunoblot.

To examine if combination of rapamycin with PLK1 inhibitors confers additional reduction of clonogenic survival, compared to rapamycin alone, we co-treated 621-101 cells with inhibitors for 72 h. Because BI-2536 appears more potent than volasertib to inhibit 621-101 cells, we treated cells with 1 and 3 nM BI-2536, and 10 and 30 nM volasertib. All single-drug treatments resulted in a statistically significant inhibition of clonogenic survival, compared to DMSO. For all concentrations tested, combination of volasertib with rapamycin significantly decreased clonogenic survival, compared to the same-dose rapamycin-only treated cells (**Figure 1C**, and **Supplemental Table 3**). Combination of BI-2536 with rapamycin did not significantly decrease clonogenic survival, compared to rapamycin-only treated cells.

Because prolonged PLK1 inhibition leads to cell death, we tested whether BI-2536, volasertib, or their combination with rapamycin induce apoptosis in 621-101 cells. Both BI-2536 and volasertib induced significant cleavage of poly (ADP-ribose) polymerase (PARP) and caspase 3, at 24 and 48 h of treatment (**Figure 1D**). Consistent with previous findings and with the notion that rapalogs are cytostatic rather than cytotoxic, rapamycin did not induce cleavage of either PARP or caspase 3. As expected, rapamycin induced phosphorylation of AKT at S473 (**Figure 1D**), a mTORC2-dependent phosphorylation event due to inhibitory phosphorylation of rictor by p70S6K (14). S473-AKT phosphorylation was attenuated, but not completely inhibited, when cells were treated for 24 h with BI-2536 or volasertib in the presence of rapamycin, but this attenuation was not observed at the 48 h timepoint. The levels of the autophagosome cargo protein p62/SQSTM1 were significantly decreased in rapamycin-treated cells, as previously shown (20, 22). Co-treatment with rapamycin and PLK1 inhibitors (BI-2536 or volasertib) attenuated this rapamycin-induced decrease of p62/SQSTM1 at the 24 h but not at the 48 h timepoint. These data are consistent with our previous observations that BI-2536 increases p62/SQSTM1 in TSC2-deficient ELT3 and 621-101 cells (20). Collectively, our data suggest that in TSC2-deficient human angiomyolipoma cells the combined PLK1 and mTORC1 inhibition by volasertib and rapamycin significantly decreases cell viability and clonogenic survival, induces apoptosis, and attenuates pro-survival AKT and p62/SQSTM1 signaling, compared to mTORC1-only inhibition.

### Volasertib decreases short-term lung colonization by TSC2-deficient ELT3 cells

LAM is caused by infiltration of the lung parenchyma with TSC2-deficient cells, probably through a mechanism involving metastatic dissemination from a distant site (23). We previously demonstrated that ELT3 cells, which are derived from an Eker rat uterine leiomyoma (24), metastasize to the lungs after subcutaneous or intravenous injection in SCID mice and that rapamycin decreases lung colonization (25). To test whether PLK1 inhibition can decrease experimental lung metastasis by ELT3 cells, we pre-treated SCID mice for two days with rapamycin (3 mg/kg ip), volasertib (25 mg/kg po), combination of volasertib and rapamycin, or vehicle control.

Luciferase-expressing ELT3 cells were then injected into the tail-vein of mice, and bioluminescence was measured at baseline and at 6 h and 24 h after inoculation (**Figure 2A, Supplemental Figure S2**, and **Supplemental Table 4**). Twenty-four (24) hours after cell injection, volasertib significantly decreased lung metastasis as demonstrated by the reduction in normalized photon flux, compared to the control group (*P* = 0.0003, **Figure 2?**, and **Supplemental Tables 4** and **5**). Volasertib also caused a significant inhibition of normalized photon flux compared to rapamycin (*P* = 0.0414). In agreement with past studies (25), rapamycin also caused a significant decrease in lung metastasis compared to control (*P* = 0.018). Six (6) hours after cell injection, no statistically significant differences have been observed between treatment groups (**Supplemental Table 6**). Collectively, our data indicate that volasertib potently inhibits ELT3 cell migration to the lungs.

**Figure 2.**
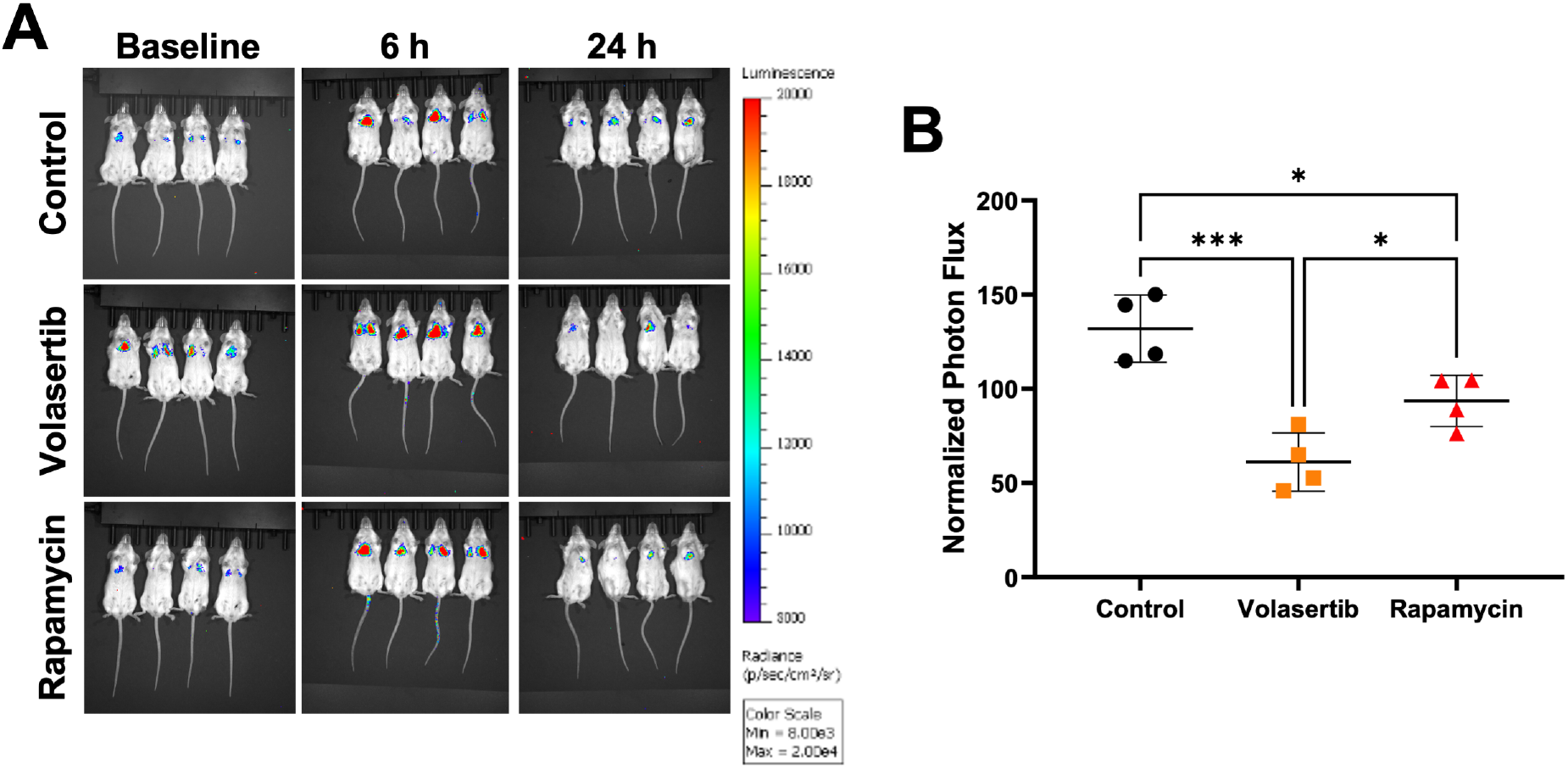
Volsaertib decreases short-term lung colonization by ELT3 cells. Eight-week-old female SCID mice were ovariectomized, supplemented with 17β-estradiol in drinking water (500 nM), and then pre-treated every 24 h for two days with vehicle control, rapamycin (3 mg/kg ip), or volasertib (25 mg/kg po). Mice were then inoculated by tail vein injection with luciferase-expressing ELT3 cells (ERL4) and underwent bioluminescence imaging at baseline, and at 6 h and 24 h after cell inoculation. **(A)** Raw bioluminescence images of mice. **(B)** Normalized photon flux [(total photon flux / group baseline average) x100] for the 24 h timepoint. Error bars are SD. Asterisks indicate statistical significance compared to the control group, or between the volasertib and rapamycin groups, based on one-way ANOVA.

### Volasertib inhibits the growth of ELT3-derived xenograft tumors

Volasertib has been shown to inhibit the formation of subcutaneous tumors derived from various cancer cell lines, including colon carcinoma, non-small cell lung cancer, rhabdomyosarcoma, neuroblastoma, hepatoblastoma, Non-Hodgkin’s Lymphoma, hepatocellular carcinoma, and acute myeloid leukemia (19, 26). ELT3 cells form subcutaneous tumors in immunodeficient mice (25) and have been used extensively in preclinical studies of experimental compounds for TSC and LAM research. Since volasertib decreased ELT3 lung metastasis, we treated SCID mice bearing subcutaneous ELT3 tumors with volasertib (25 mg/kg po twice a week on two consecutive days), rapamycin (3 mg/kg ip three times a week), or combination of volasertib and rapamycin. Compared to control, volasertib significantly inhibited tumors from day 4 after start of treatment (**Figure 3A, Supplemental Tables 7 and 8**; *P* < 0.05 two-way ANOVA). This volasertib-induced reduction in tumor size was sustained until the end of treatment at day 28. As previously reported, rapamycin treatment induced a very significant inhibition of ELT3-derived tumors, compared to control, starting at day 2 post treatment (**Figure 3A, Supplemental Tables 7 and 8)**, and reduced the tumors to non-palpable after a week of treatment. Due to the very robust inhibition of tumor growth by rapamycin, combination treatment did not have a significant difference from rapamycin-only. Taken together, our results show that PLK1 inhibition by volasertib significantly hinders the growth of TSC2-null xenograft tumors.

**Figure 3.**
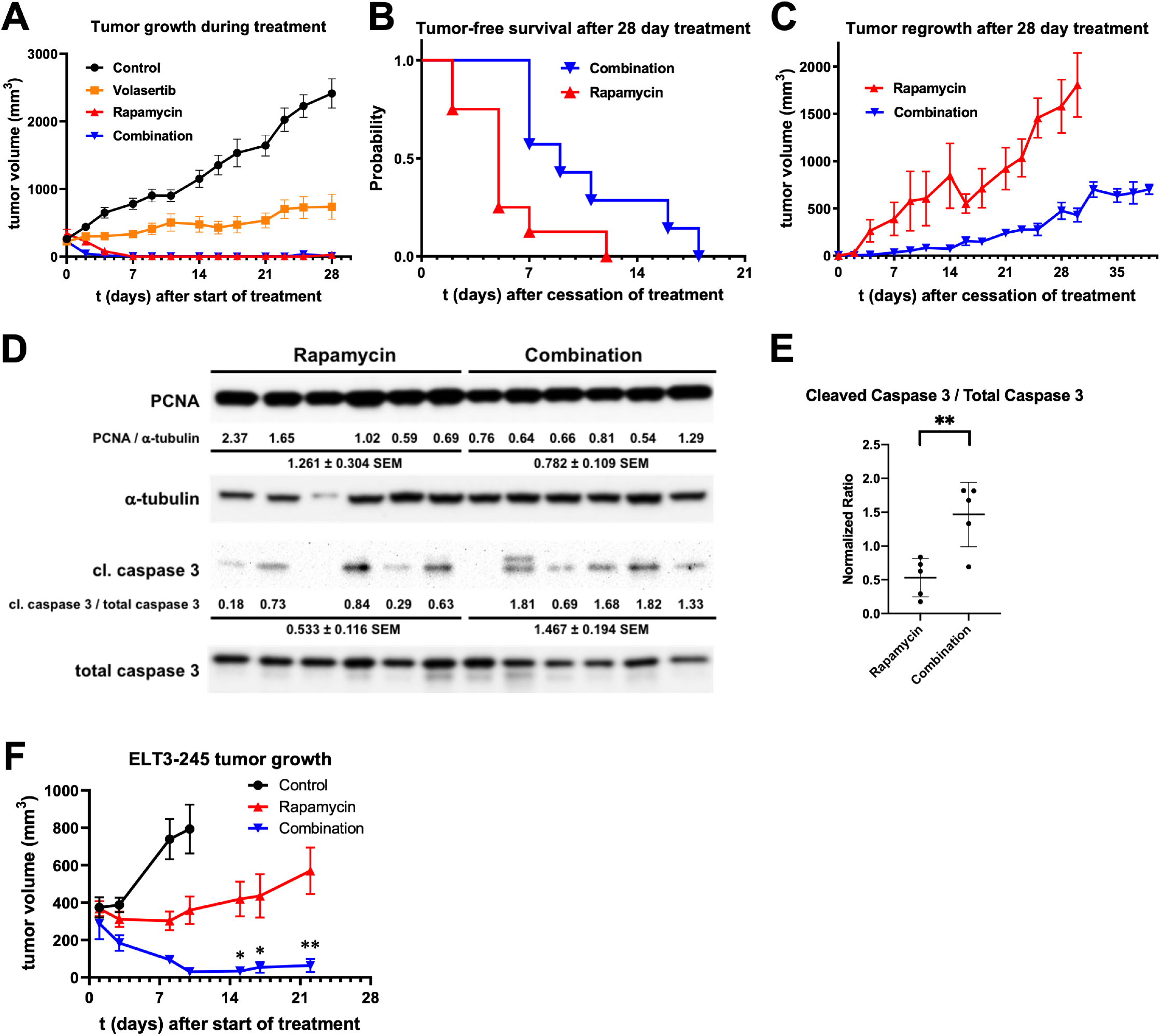
Combination of volasertib and rapamycin inhibits the regrowth of ELT3-derived xenograft tumors. **(A)** Average ELT3 xenograft tumor volume (mm^3^, *y* axis) during a 28-day treatment (*x* axis) with volasertib (25 mg/kg po twice a week on two consecutive days, orange squares, n=24), rapamycin (3 mg/kg ip, three times a week, red upward triangles, n=18), and combination of volasertib and rapamycin (blue downward triangles, n=21), compared to vehicle control (closed circles, n=27). Error bars are SEM. **(B)** Tumor-free survival of mice after cessation of a 28-day treatment with rapamycin (median survival 5 days, n=9, red upward triangles) or combination of volasertib and rapamycin (median survival 9 days, n=8, blue downward triangles). Log-rank (Mantel-Cox) test P=0.0216 *, Gehal-Breslow-Wilcoxon test P=0.0094 **. **(C)** Average ELT3 xenograft tumor volume (mm^3^, *y* axis) after cessation of a 28-day treatment with rapamycin (red closed upward triangles, red solid line) or combination of volasertib and rapamycin (blue closed downward triangles, blue solid line). Error bars are SEM. (**D**) Regrowth tumor lysates from rapamycin- and combination-treated mice (n=6 per group) were resolved by PAGE and immunoblotted with the corresponding antibodies. The PCNA / α-tubulin and cleaved caspase 3 / total caspase 3 ratios for each sample (normalized to the average ratio for all samples) are shown under the PCNA and cleaved caspase 3 immunoblots, respectively. (**E**) Band densitometric analysis of Cleaved caspase 3 / Total Caspase 3 ratio from the immunoblot presented in Figure 3D. Rapamycin: 0.533±0.116 SEM, n=5; Combination 1.467±0.194 SEM, n=5; P=0.0055 * unpaired T-test. Error bars are SD. (**F**) SCID mice were inoculated with ELT3-245 cells and treated 3 times per week with vehicle (closed circles, black line), rapamycin (3 mg/kg ip three times a week, red upward triangles, red line), or combination of volasertib (17 mg/kg ip three times a week) and rapamycin (3 mg/kg ip three times a week). Error bars are SEM. Asterisks indicate statistical significance between the rapamycin-only and the combination groups, based on two-way ANOVA.

### Combination of volasertib and rapamycin reduces ELT3 tumor regrowth after discontinuation of treatment

To test the effect of combination treatment on the regrowth of ELT3 subcutaneous xenograft tumors after discontinuation of treatment, a subset of mice that were treated for 28 days with rapamycin or combination of volasertib and rapamycin were left untreated and the regrowth of tumors was monitored. Mice that had been treated for 28 days with combination of volasertib and rapamycin had significantly increased median tumor-free survival, compared to mice that received rapamycin monotherapy (**Figure 3B**). All tumors from combination treated mice beyond day 21 after cessation of treatment were significantly smaller compared to regrowth tumors from rapamycin only treated mice (**Figure 3C** and **Supplemental Table 9**). Similarly, mice that had been treated with combination for 14 days had a slower tumor regrowth (**Supplemental Figure S3A** and **Supplemental Table 10**) and increased tumor-free survival (**Supplemental Figure S3B**), compared to those that had been treated with rapamycin-only for 14 days. Comparing tumor-free survival after cessation of treatment, there were no differences between 14-day and 28-day rapamycin treatment (**Supplemental Figure S4A**) and between 14-day and 28-day combination treatment (**Supplemental Figure S4B**). Of note, the regrowth tumors from 14-day combination treated mice were smaller, compared to those from 28-day rapamycin treated mice (**Supplemental Figure S4C** and **Supplemental Table 11**) although there was no statistically significant difference. The 14-day combination treated mice had increased tumor-free survival, compared to the 28-day rapamycin treated mice (**Supplemental Figure S4D**) without statistically significant difference. Collectively, these data demonstrate that rapamycin and volasertib combination treatment significantly retards TSC2-null tumor regrowth, compared to rapamycin monotherapy.

### Regrowth tumors from combination-treated mice have sustained apoptosis after discontinuation of treatment

Since tumors from 28-day combination-treated mice regrew slower than those from 28-day rapamycin-only treated mice, we sought to evaluate the effects of combination treatment on proliferation and apoptosis. We analyzed age-matched regrowth tumors from each of the two treatment groups at approximately 35 days after cessation of treatment (**Supplemental Table 12**). Since tumors from the combination treated mice regrew slower, compared to tumors from rapamycin treated mice, and this size difference could affect changes in proliferation and apoptosis rates, we tried to match tumor volume between the two treatment groups to the extent possible. Immunoblotting showed that regrowth tumors from combination treated mice had significantly increased cleaved caspase 3 / total caspase 3 ratio, compared to regrowth tumors from rapamycin treated mice (**Figures 3D** and **3E**), suggesting increased apoptosis in regrowth tumors from combination-treated mice. Lower PCNA levels were observed in the regrowth tumors from combination treated mice, compared to rapamycin treated, although this difference was not statistically significant (**Supplemental Figure S5**).

### Combination treatment decreases the growth of rapamycin-refractory ELT3-245 xenograft tumors

We previously reported the development of an ELT3 derivative cell line, termed ELT3-245, with increased tumorigenicity in SCID mice, compared to the parental ELT3 cells (27). ELT3-245-derived xenograft tumors are refractory to rapamycin treatment. Given the effect of volasertib and rapamycin combination to decrease ELT3 tumor regrowth, we treated ELT3-245 tumor-bearing mice with rapamycin, or combination of volasertib and rapamycin. Combination treatment significantly inhibited ELT3-245 tumor growth 10 days after start of treatment, and this was sustained through day 22 (**Figure 3F** and **Supplemental Table T13**). As we previously reported (27), rapamycin alone did not inhibit ELT3-245 xenograft tumor growth.

### Regrowth tumors from combination-treated mice have differential expression of type I interferon and interleukin signaling pathways after discontinuation of treatment

To identify genes that are differentially expressed in the regrowth tumors from combination treated mice, compared to rapamycin, we used RNA from the tumors described in Supplemental Table 12 and performed genome-wide gene expression microarray analyses. Out of the 19,395 unique genes in the array, 105 genes were differentially regulated in the regrowth tumors from combination treated mice, compared to those from rapamycin treated mice (70 genes upregulated, 35 genes downregulated, P<0.05, FDR<0.05, **Supplemental Table 14**). Integrated Pathway Analysis (IPA) identified “Interferon Signaling”, “Pathogenesis of Multiple Sclerosis”, “Role of JAK2 in Hormone-like Cytokine Signaling”, “IL-9 Signaling”, and “Role of JAK1 and JAK3 in Cytokine Signaling” as the top five differentially regulated pathways. Of the 105 differentially expressed genes, ten (10) were included in the top five pathways: *Cxcl10, Cxcl11, Ifit3, Il7, Isg15, Mx1, Oas1, Socs2, Stat5a* and *Stat5b* (**Supplemental Table 15** and **Supplemental Figure S6**).

We next threaded the list of 105 differentially expressed genes in the ToppGene application (28, 29) using the human ortholog gene names. Functional gene enrichment analysis showed that the top Gene Ontology (GO) Biological Process annotation was GO:0035455 “response to interferon-alpha”, and the top five (5) pathways were “Interferon alpha/beta signaling”, “Immune response to tuberculosis”, “Cytokine Signaling in Immune system”, “IL-7 Signal Transduction”, and “Measles”. These six (6) annotations included 17 of the 105 differentially expressed genes (**Figure 4A, Table 1**, and **Supplemental Table 16**). We threaded these 17 genes into the STRING database (30) to obtain a network of functional and physical interactions (**Figure 4B**). This network comprises two distinct sub-networks: one related to interferon signaling and the other to interleukin signaling. Gene regulation withing the two sub-networks is opposite, with the genes in the interferon signaling sub-network being downregulated and the genes in the interleukin signaling sub-network being mostly upregulated. To validate these gene expression data, we performed independent RT-qPCR assays for 6 of these 17 genes and confirmed their differential expression in regrowth tumors from combination-vs. rapamycin-treated mice (**Figure 4B** and **Table 1**).

**Figure 4.**
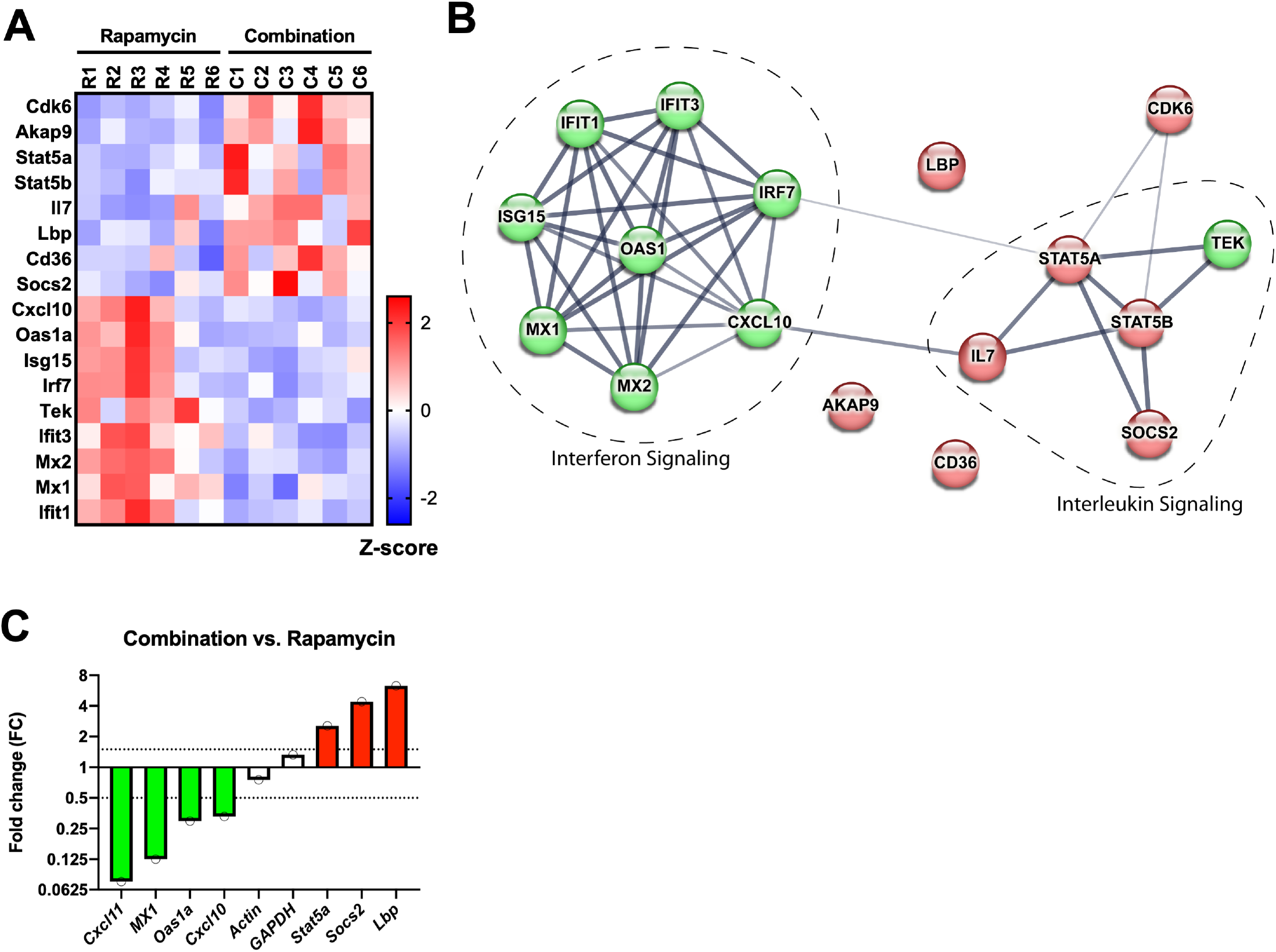
Regrowth tumors of combination-treated mice have differential expression of type I interferon and interleukin signaling pathways. Regrowth tumors at approximately 35 days after discontinuation of rapamycin-only or combination treatment were analyzed by global gene expression arrays **(A)** Gene expression heatmap of 17 type I interferon and interleukin signaling pathways, sorted by row P value. (B) Functional and physical interaction network of the 17 differentially expressed genes from panel A. Nodes in red are upregulated genes. in regrowth tumors from combination treated mice, compared to rapamycin. Nodes in green are downregulated genes. Connecting lines between nodes indicate interactions. High-confidence interactions are shown with more prominent lines. (C) RT-qPCR validation of differentially expressed genes in regrowth tumors from combination-treated mice, compared to regrowth tumors from rapamycin-treated mice. Thresholds are indicated with the horizontal dashed lines intersecting the y axis at FC 0.5 and 1.5. Genes with green bars are downregulated (FC < 0.5), genes with red bars are upregulated (FC > 1.5). *Actin* and *Gapdh* were used as housekeeping control genes (white bars, 0.5<FC<1.5).

**Table 1.** The 17 differentially expressed genes in top annotations from ToppGene gene enrichment. Fold-change and regulation is from comparing regrowth xenograft tumors from combination-treated vs. rapamycin-treated mice (n=6 per group). For select genes, RT-qPCR validation was performed.

|  | ARRAY DATA |  | RT-qPCR DATA |  |  |
| --- | --- | --- | --- | --- | --- |
|  | FC | Regulation | FC | Fold Regulation | Regulation |
| *Lbp* | 11.829 | UP | 6.326 | 6.326 | UP |
| *Socs2* | 5.001 | UP | 4.405 | 4.405 | UP |
| *Stat5a* | 4.206 | UP | 2.556 | 2.556 | UP |
| *Akap9* | 4.039 | UP |  |  |  |
| *Il7* | 3.253 | UP |  |  |  |
| *Stat5b* | 3.190 | UP |  |  |  |
| *Cd36* | 2.626 | UP |  |  |  |
| *Cdk6* | 2.544 | UP |  |  |  |
| *Tek* | 2.407 | DOWN |  |  |  |
| *Ifit3* | 2.550 | DOWN |  |  |  |
| *Oas1a* | 2.744 | DOWN | 0.296 | 3.382 | DOWN |
| *Isg15* | 3.254 | DOWN |  |  |  |
| *Mx1* | 3.343 | DOWN | 0.125 | 7.985 | DOWN |
| *Ifit1* | 3.390 | DOWN |  |  |  |
| *Mx2* | 3.558 | DOWN |  |  |  |
| *Irf7* | 3.764 | DOWN |  |  |  |
| *Cxcl10* | 3.835 | DOWN | 0.329 | 3.044 | DOWN |
| *Cxcl11* * | 5.569 | DOWN | 0.075 | 13.284 | DOWN |

\* *Cxcl11* is not among the 17 genes in the top annotations.

## DISCUSSION

Although aberrant PI3K/mTOR is the main signaling defect in TSC and LAM lesions, the severity, progression, and outcomes of these lesions are highly inconsistent and depend on variables such as age of onset and, in the case of LAM, on gender. Although most individuals affected by TSC and LAM benefit from the use of rapalogs, a small subset of patients do not respond to this treatment and their tumors do not regress or, even worse, continuously grow despite being on treatment. Based on our previous works offering a novel insight on the potential of PLK1 as a targeted therapy for TSC and LAM (5, 6, 20), we addressed whether exploiting this with a pharmacological approach *in vivo* would offer a benefit over traditional rapalog regimens. We found that, similar to the PLK1 inhibitor BI-2536, volasertib decreased the viability and clonogenic survival and induced apoptosis of the LAM patient-derived angiomyolipoma cell line 621-101. In combination with rapamycin, volasertib significantly inhibited the clonogenic survival of 621-101, decreased the rapamycin-induced phosphorylation of S473-AKT and attenuated the degradation of p62, consistent with our previous findings of BI-2536 effects on TSC1- and TSC2-deficient cells (20). Sporadic pulmonary LAM affects almost exclusively women (31), is caused primarily by *TSC2* mutations (4) and is associated with lung parenchyma infiltration by TSC2-deficient smooth muscle-like cells (23). Previously we demonstrated that the TSC2-deficient ELT3 cells migrate to the lungs of SCID mice, and that estrogen increases their capacity to colonize the lungs (25). Using luciferase-expressing ELT3 cells, we showed that volasertib decreases lung colonization. Collectively, the above data suggest that inhibition of PLK1 impedes the survival and metastatic potential of TSC2-deficient cells.

ELT3 cells are tumorigenic in immunodeficient mice and have been extensively used in preclinical studies for new therapeutics for TSC and LAM. We sought to test the efficacy of volasertib to inhibit tumorigenesis of TSC2-deficient cells in the ELT3 xenograft mouse model. During a 28-day course of treatment, volasertib alone significantly inhibited the growth of ELT3 tumors, compared to control-treated mice. As expected, rapamycin almost completely inhibited tumor growth between 4 and 7 days of treatment.

Combination of volasertib and rapamycin seemed to be superior to inhibit ELT3 tumors, compared to rapamycin alone, at very early time-points of treatment. Due to the very significant tumor inhibition by rapamycin, it was not feasible to determine the contribution of volasertib to a potential synergistic effect. More importantly, when studying tumors derived from the rapamycin-refractory ELT3-245 cells (27), combination of volasertib and rapamycin caused an almost complete inhibition of tumor growth.

Since discontinuation of rapalog treatment in TSC and LAM causes regrowth of lesions (10, 32), we wanted to study the effects of combination treatment withdrawal on ELT3 tumors. ?ice treated with combination of volasertib and rapamycin had a significantly slower tumor regrowth rate after discontinuation of treatment, compared to rapamycin only. This difference was sustained until one month after discontinuation of treatment, when the regrowth tumors from rapamycin-treated mice reached secondary endpoint criteria. We obtained similar results with regrowth tumors from mice treated with combination or rapamycin for only 14 days, although the regrowth rate in the latter groups was accelerated compared to the 28-day treatment groups. Tumor-free survival after discontinuation of treatment was also significantly improved in combination-treated mice, compared to rapamycin only. Notably, a 14-day combination treatment resulted in a better inhibition of tumor regrowth, compared to a 28-day rapamycin treatment. Although these differences did not reach statistical significance, most likely due to the small number of regrowth tumors in the 14-day combination treatment group (*n* = 4), we believe that they warrant further investigation.

Approximately one month after discontinuation of treatment, regrowth tumors from the combination treated mice showed evidence of increased apoptosis, compared to regrowth tumors from the rapamycin group. Using global gene expression profiling, we found that genes involved in type I interferon signaling were downregulated in regrowth tumors from combination treated mice, compared to rapamycin only. Type I interferons activate the interleukin/JAK/STAT signaling pathway to upregulate the transcripts of interferon stimulated genes (33), including *Oas1, Mx1*, and *Isg15* that were downregulated in the regrowth tumors from the combination-treated group. Opposing to the downregulation of type I interferon signaling genes in regrowth tumors from combination-treated mice, genes in the interleukin/JAK/STAT signaling pathway were overexpressed. Type I interferons have been associated with favorable outcomes in cancer therapy, as they play an important role in tumor immunosurveillance (34). It was previously shown that an *IFNG* (interferon-γ) high-expression allele is associated with lower incidence of renal angiomyolipomas in TSC (35), and that combination of interferon-γ with the rapalog CCI-779 was more effective to decrease TSC mouse lesions, compared to CCI-779 alone (36). Finally, rapamycin suppresses interferon production by dendritic cells (37), whereas the Rheb inhibitor S-trans,trans-farnesylthiosalicylic acid results in the differential regulation of type I interferon signaling (38).

Mining publicly available protein-protein interaction databases, we found that PLK1 associates with interferon signaling through a physical interaction with the mitochondrial antiviral signaling adapter protein MAVS (39). MAVS, through the upregulation of TNF Receptor-Associated Factors (TRAFs) and Interferon Regulator Factors (IRFs) enhances anti-viral innate immunity and the production of type I interferons (40). By binding MAVS, PLK1 inhibits the activation of TRAFs and IRFs and decreases interferon production, whereas depletion of PLK1 enhances production of type I interferons (39). Interestingly, MAVS forms prion-like structures to sustain long-term anti-viral immune responses (41). We speculate that PLK1 inhibitor treatment not only initiates apoptotic events and anti-autophagic signaling to decrease the survival of TSC2-deficient cells in the tumors but may augment long-term tumor remission by facilitating a sustained MAVS-interferon signaling cascade.

Two preclinical studies provide evidence of apoptosis in ELT3 xenograft models during treatment. The first study used a combination of rapamycin and the autophagy inhibitor resveratrol (42), while the second study used the combination of nelfinavir, an HIV protease inhibitor with broad anti-cancer activity (43), and the proteasome inhibitor bortezomib (44). To our knowledge, our current study is the only one presenting evidence of sustained apoptosis in TSC2-deficient regrowth tumors approximately one month after discontinuation of treatment. Based on the terminal half-life for volasertib (46 h) and rapamycin (15 h) in mice (19, 45), we estimate that plasma concentrations at 35 days after discontinuation of treatment would be less than 0.001% of the initial plasma concentration for both drugs. Therefore, the increased apoptosis we observed in the regrowth tumors could not be attributed to a systemic presence of volasertib in mice. A thorough pharmacokinetic study for any potential drug-drug interaction between volasertib and rapamycin is necessary.

In conclusion, our data support that combined mTORC1 and PLK1 inhibition can potently interfere with the regrowth of TSC and LAM relevant tumors for a long time after discontinuation of treatment and with the growth of rapamycin-refractory tumors. Thus, combination of rapalogs with PLK1 inhibitors may produce faster clinical responses and sustained remissions for TSC and LAM tumors.

## MATERIALS AND METHODS

Detailed procedures are described in Supplemental Information.

### Cell cultures and treatments

621-101 (21), ELT3 (24) and ELT3-245 (27) cells were cultured in IIA complete media at 37°C and 5% CO_2_ air. Absolute counts of living and dead cells were obtained by trypan blue exclusion and automated cell counting. Rapamycin (CAS #53123-88-9), BI-2536 (CAS #755038-02-9), and volasertib (CAS #755038-65-4) were purchased from Selleck Chemicals (S1039, S1109, and S2235, respectively). Compounds were dissolved at 10 mM in dimethyl sulfoxide (DMSO, Sigma D2650) and stored at −20°C.

### Cell viability and survival assays

For viability, cells were seeded in 96-well plates, and treated with various concentrations of compounds for 48 h. For cell survival, cells were treated with compounds for 72 h, then equal number of living cells were allowed to form colonies for 14 days.

### Mouse studies

All animal studies were approved by the Institutional Animal Care and Use Committees at the University of Tennessee Health Science Center (protocol #16-166), Texas Tech University Health Sciences Center (protocol #14031), and the University of Cincinnati (protocol # TR01-15-07-22-01). Procedures were performed according to all relevant guidelines and regulations. Eight-week-old female Fox Chase SCID (CB17 SCID) mice were obtained from Taconic (CB17SC-F EF) or The Jackson Laboratory (B6.CB17-Prkdcscid/SzJ). For short-term lung colonization studies (25), ovariectomized mice were supplemented with 17β-estradiol in drinking water, pre-treated with compounds, and inoculated intravenously with luciferase-expressing ELT3 cells. Bioluminescence was measured at 1 h, 6 h and 24 h post inoculation. For xenograft mouse models, mice were inoculated with cells bilaterally by subcutaneous injection, and tumor volume was calculated based on caliper measurements. Mice were sacrificed when primary or secondary endpoint criteria were met.

### Protein analyses

Cells were lysed in PTY buffer. Tumor samples were homogenized in TPER buffer. Proteins were resolved in polyacrylamide gels, transferred on PVDF membranes, and immunoblotted with primary and secondary antibodies.

### Gene expression studies

Tumor RNA was used for genome-wide expression arrays and validation of data by RT-qPCR.

### Statistics and Graphing

Except for global gene expression microarray profiling, all other *in vitro* studies were performed at least three times independently. The two-sample unpaired Student’s T-test was used for comparisons between treatment groups. Statistical analysis (T-test and ANOVA), descriptive statistics (arithmetic mean, standard deviation (SD), Standard Error of the Mean (SEM)), and graphing were performed with Prism 9 (version 9.4.1, GraphPad Software, LLC). Statistical significance indicators are * for P < 0.05, ** for P < 0.01, *** for P < 0.001, and **** for P < 0.0001.

### Digital image acquisition and editing

Digitally captured images or scanned images of X-ray films of immunoblots were saved in uncompressed TIFF format. Digital images were edited using Adobe Photoshop CC (Release 2017.0.1). For presentation of final figures, images were cropped, and intensity levels were modified throughout the entire cropped section.

## Acknowledgements

We thank Mr. Rahul Mavinkurve for technical work, and Mr. Lorne Rose (Molecular Resource Center, UTHSC) for gene expression microarray assays. This work was partially funded by the University of Pennsylvania Orphan Disease Center grant number MDBR-15-103-LAM (to AA) and by NIH-R01HL138481 (to JJY).

