## Supplemental Data for "Combination of Volasertib and Rapamycin Inhibits the Regrowth of TSC2-Deficient Tumors"

Supplemental Tables 1-16 can be found in the Excel file labeled “Supplemental_Tables.xlsx”.

**Supplemental Figure S1.** CumpuSyn output for dose-effect curve for volasertib (Vol, red line, red open squares) and BI-2536 (blue line, blue open circles) after 48 h treatment of 621-101 cells. *x* axis: drug concentration (nM). *y* axis: effect (Fa) on cell viability.

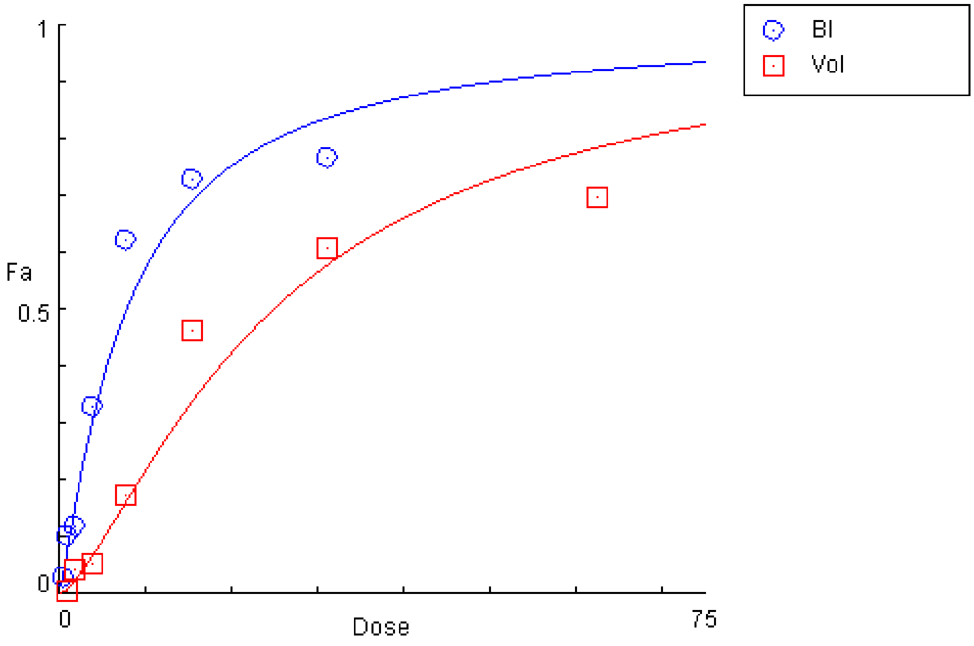

**Supplemental Figure S2. (A)** Total photon flux (photons per second [p/s]) after inoculation with ERL4 (luciferase-expressing ELT3) cells. **(B)** Normalized photon flux (NPF, normalized to the average total photon flux for the baseline measurement in each pre-treatment group) of data shown in (A) and in Supplemental Table S3. Vehicle (control, closed circles), volasertib (orange squares), rapamycin (red upward triangles), or combination (blue downward triangles). Error bars for both A and B are SD.

**A**

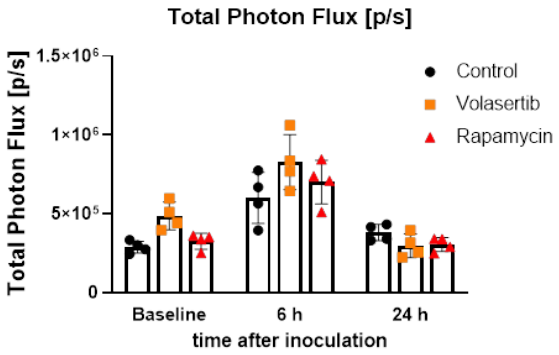


**B**

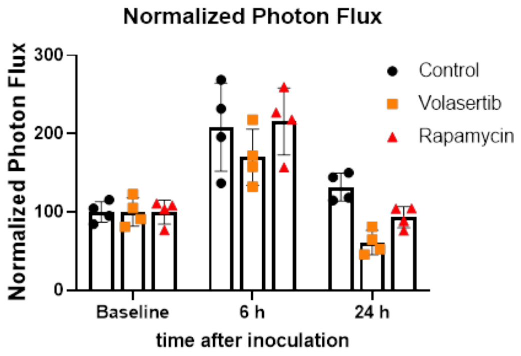

**Supplemental Figure S3. (A)** Comparison of ELT3 xenograft tumor regrowth after discontinuation of treatment between mice treated with rapamycin for 14 days (n=6, red closed upward triangles, red dotted line) and combination for 14 days (n=4, blue closed downward triangles, blue dotted line). Error bars are SEM. (**B**) Tumor-free survival after discontinuation of 14-day rapamycin treatment (median survival 3.5 days, n=6, red closed upward triangles, red dotted line) compared to 14-day combination treatment (median survival 10.5 days, n=4, blue closed downward triangles, blue dotted line). Median Log-rank (Mantel-Cox) test P=0.0339 *, Gehan-Breslow-Wilcoxon test P=0.0964 n/s.

**A**

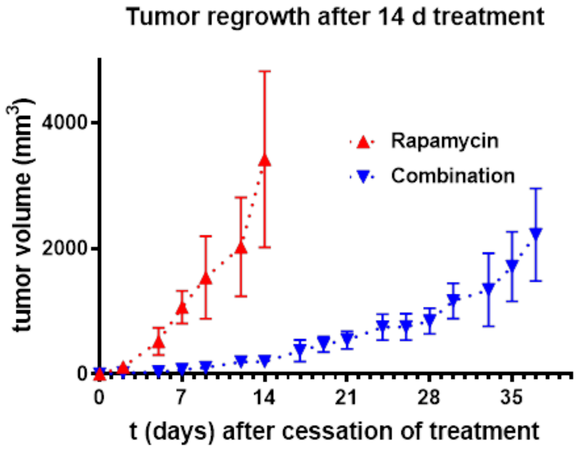


**B**

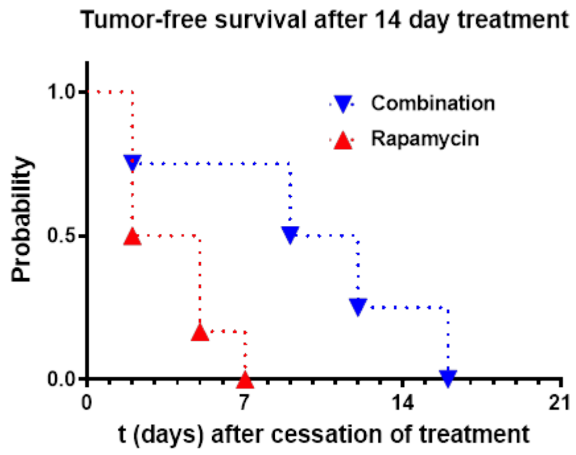

**Supplemental Figure S4. (A)** Tumor-free survival after discontinuation of treatment in mice treated with rapamycin for 14 days (n=6, red open upward triangles, red dotted line) compared to mice treated with rapamycin for 28 days (n=8, red closed upward triangles, red solid line). Median Log-rank (Mantel-Cox) test P=0.3421 n/s, Gehan-Breslow-Wilcoxon test P=0.3681 n/s. **(B)** Tumor-free survival after discontinuation of treatment in mice treated with combination for 14 days (n=4, blue open downward triangles, blue dotted line) compared to mice treated with combination or 28 days (n=8, blue closed downward triangles, blue solid line). Median Log-rank (Mantel-Cox) test P=0.8179 n/s, Gehan-Breslow-Wilcoxon test P>0.9999 n/s. (**C**) Comparison of ELT3 xenograft tumor regrowth after discontinuation of treatment between mice treated with rapamycin for 28 days (n=8, red closed upward triangles, red solid line) and combination for 14 days (n=4, blue closed downward triangles, blue dotted line). Error bars are SEM. (**D**) Tumor-free survival after discontinuation of treatment in mice treated with rapamycin for 28 days (n=8, red closed upward triangles, red solid line) compared to mice treated with combination or 14 days (n=4, blue open downward triangles, blue dotted line). Median Log-rank (Mantel-Cox) test P=0.1275 n/s, Gehan-Breslow-Wilcoxon test P=0.1960 n/s.

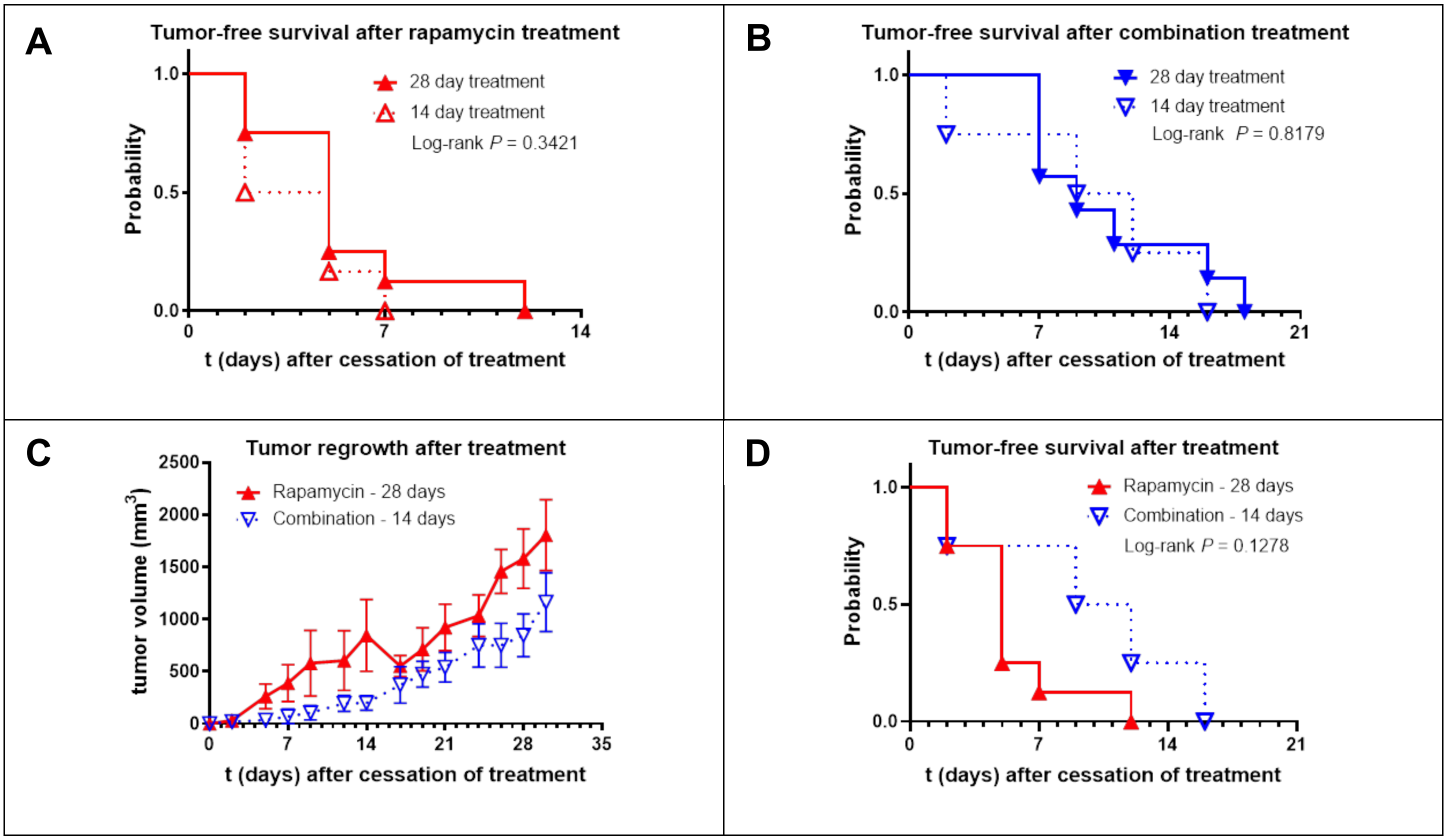

**Supplemental Figure S5.** Band densitometric analysis of PCNA / α-tubulin ratio presented in Figure 3D. Rapamycin: 1.261±0.304 SEM, n=5; Combination 0.782±0.109 SEM, n=6; P=0.1734 n/s, unpaired T-test. Error bars are SD.

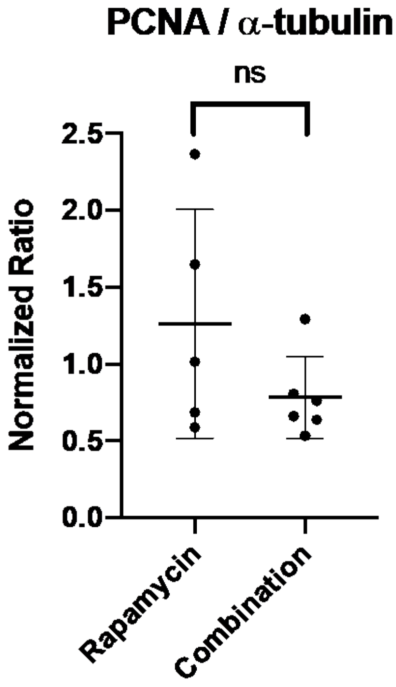

**Supplemental Figure S6.** Differentially regulated pathways in regrowth tumors from combination- vs. rapamycin-treated mice, based on gene expression array data and Ingenuity Pathway Analysis. **(A)** Interferon Signaling Pathway.

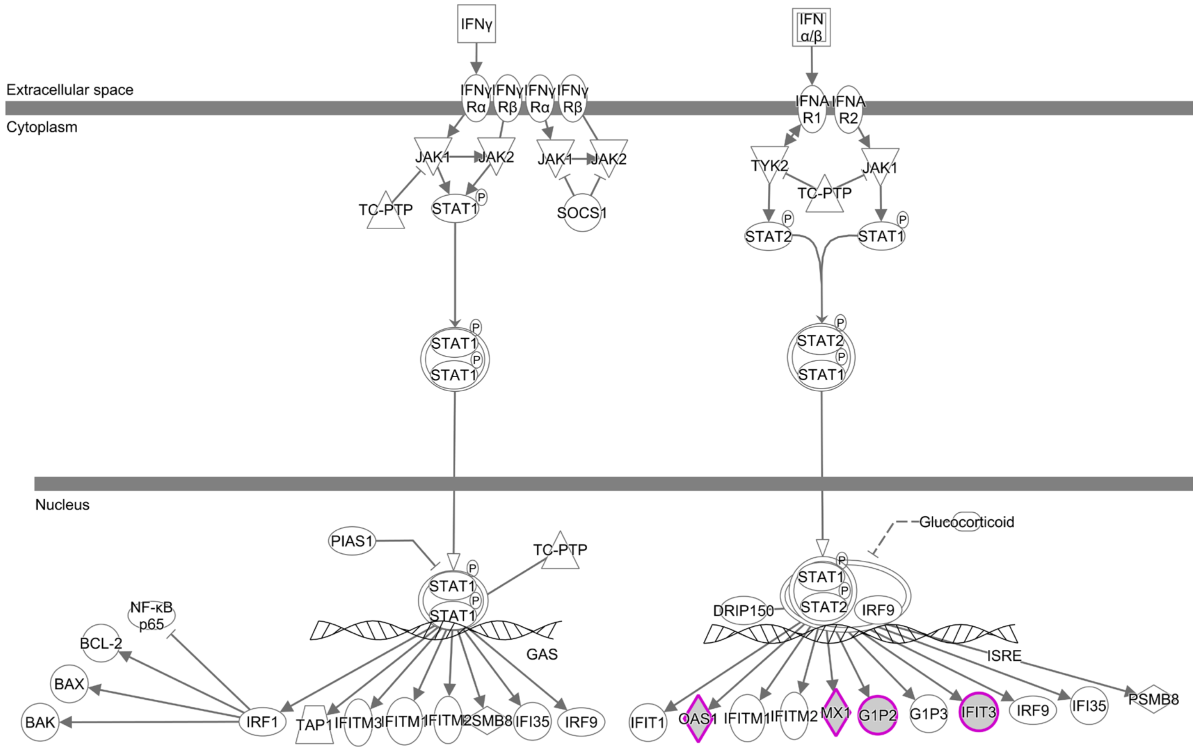


**(B)** Pathogenesis of Multiple Sclerosis Pathway.

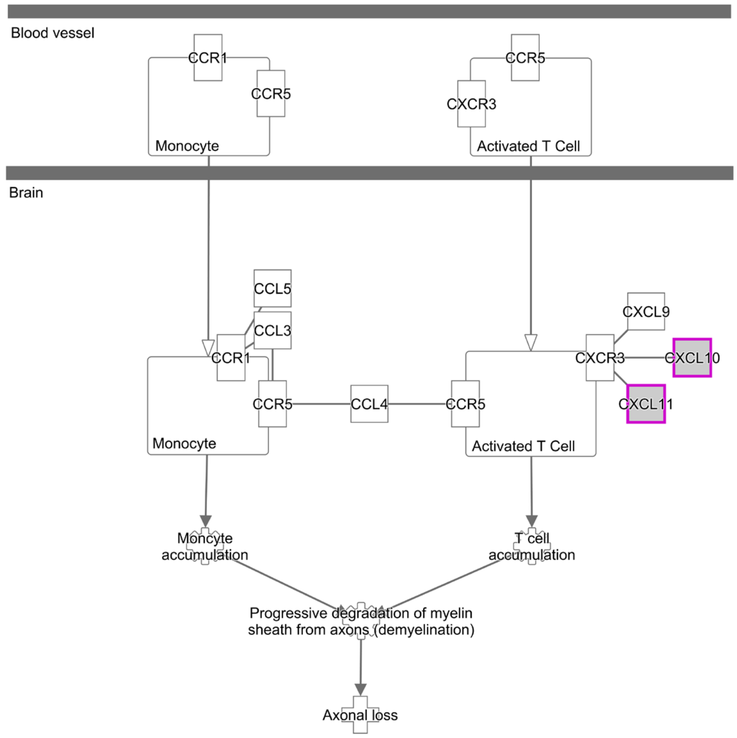

**Supplemental Figure S6.** (continued) **(C)** Role of JAK2 in Hormone-like Cytokine Signaling Pathway.

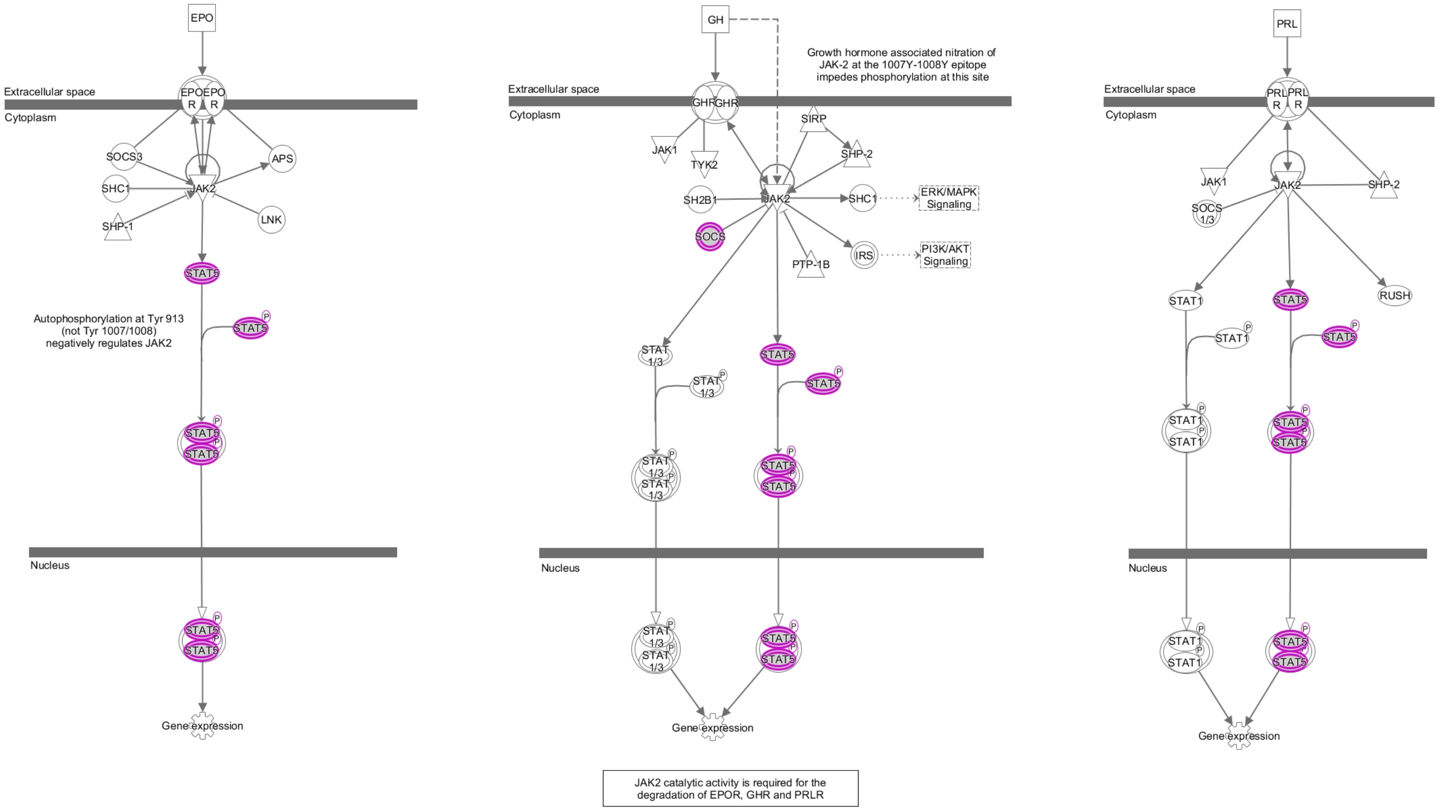

**Supplemental Figure S6.** (continued) **(D)** IL-9 Signaling Pathway.

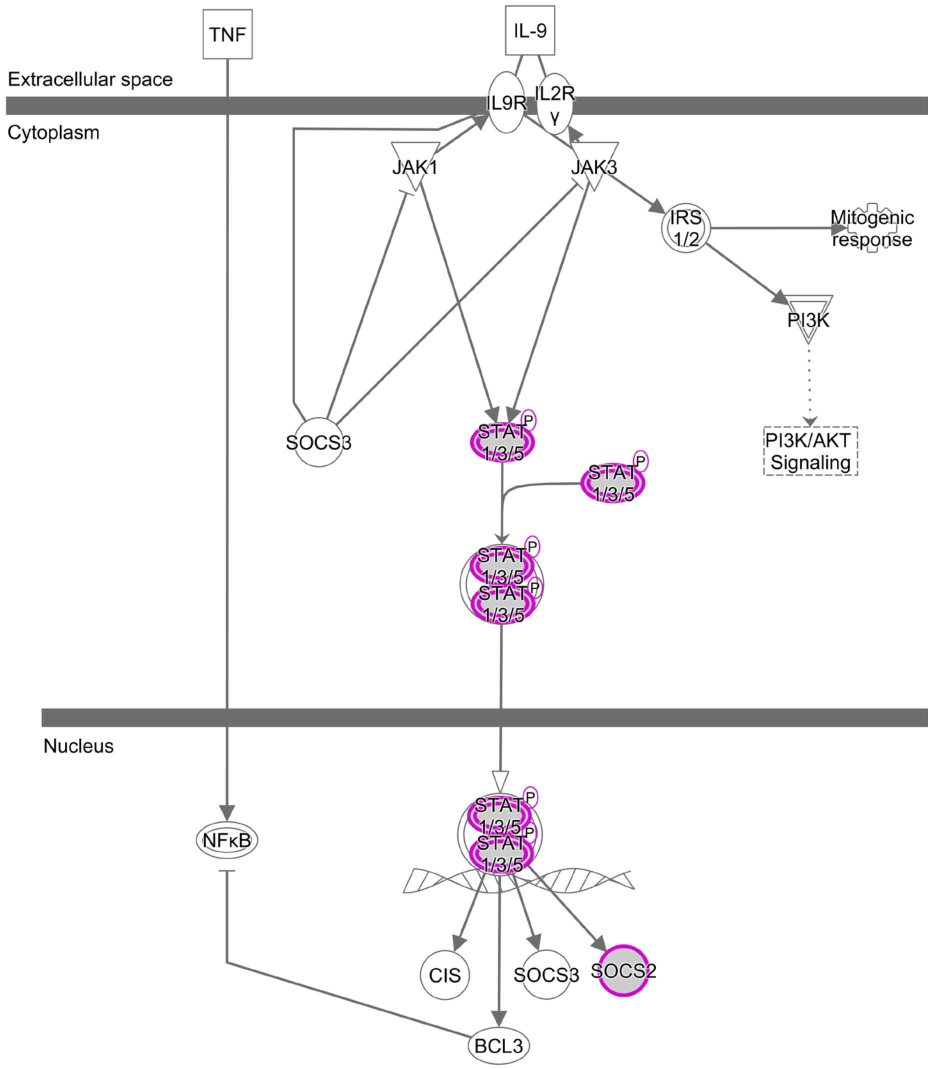

**Supplemental Figure S6.** (continued) **(E)** Role of JAK1 and JAK3 in Cytokine Signaling Pathway.

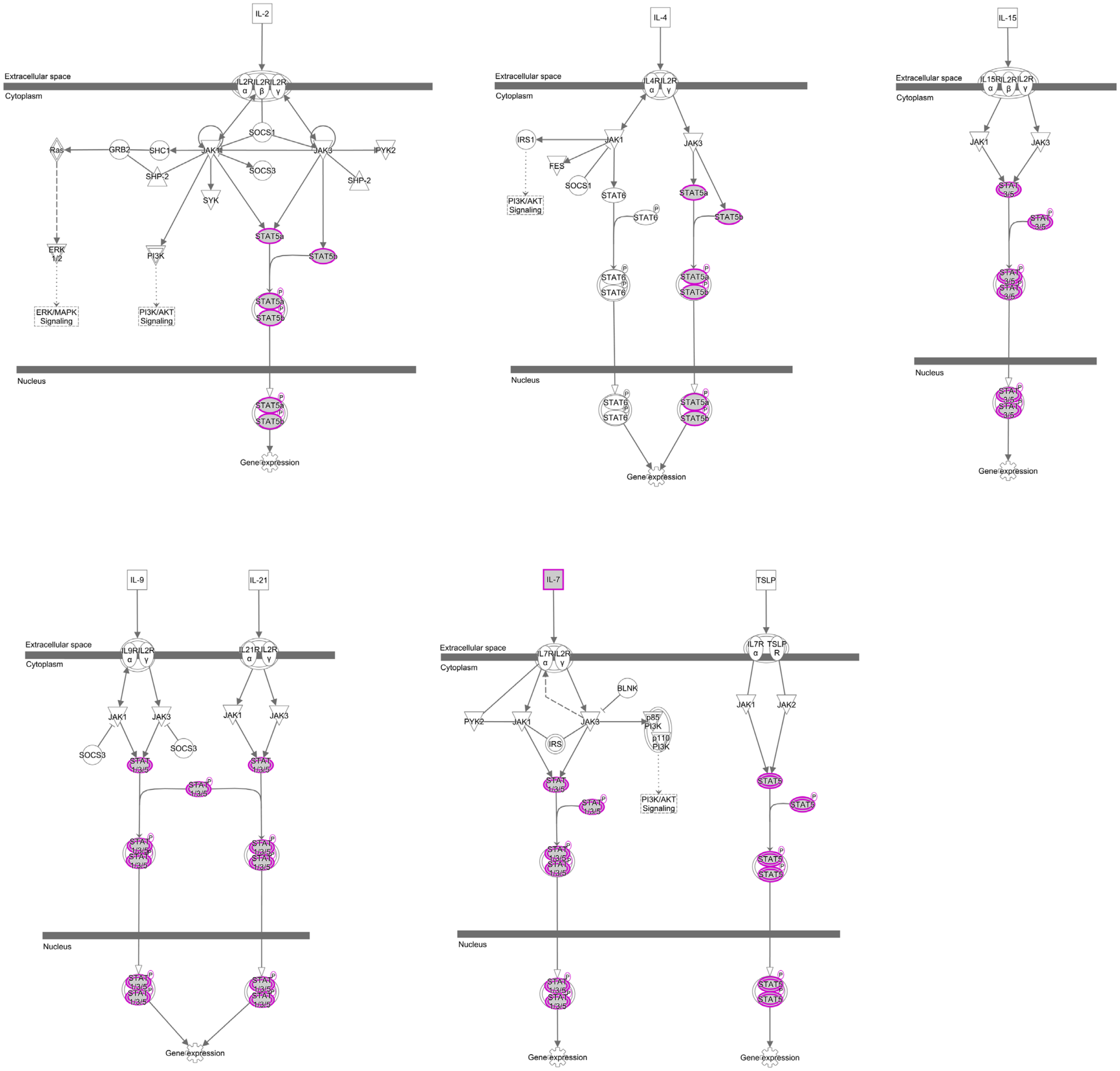
