## Supplemental Information for "Combination of Volasertib and Rapamycin Inhibits the Regrowth of TSC2-Deficient Tumors"

### SUPPLEMENTAL MATERIALS AND METHODS

**Cell Culture Media.** IIA complete media consist of DMEM/F12 50:50 without phenol red (Sigma D6434 or Corning 16405CV), 50 nM $Na_2SeO_3$ (Sigma S9133), 1.6 μM $FeSO_4$ (Fisher Scientific I146), 25 μg/ml insulin (Invitrogen A11382IJ), 0.2 μM hydrocortisone (Sigma H4001), 10 μg/ml holotransferrin (Sigma T0665), 1 nM triiodothyronine (Sigma T2752), 10 μU/ml Arg8-vasopressin (Sigma V0377), 10 nM cholesterol (Sigma C3045), 20 ng/ml epidermal growth factor (Corning 354001)], 5 mM L-glutamine (Corning 25005CI), 10% v/v fetal bovine serum (FBS, GE Healthcare Life Sciences SH3007103 or Gibco A3160501), and 100 U/ml penicillin and 100 μg/ml streptomycin (Corning 30002CI).

**Cell viability assay, and determination of GI50.** All liquid handling was performed with a Precision XS Microplate Sample Processor (Biotek) using Precision Power Automated Microplate Pipetting System software (Biotek). Each experiment was conducted at least 3 times independently, with intra-assay technical octuplicates. Cells were plated in columns 1-11 of 96-well cell culture plates ($10^2$-$10^3$ cells in 100 μl per well). Mock-treated wells (mock), containing only culture media but no cells, were added in column 12. Cells were allowed to attach for 16-24 hours. Two-fold serial dilutions of compounds (diluted in culture media) were added in columns 1-10 (25 μl per well). Culture media (with vehicle) were added in columns 11 and 12. Cells were then cultured in a 37°C 5% $CO_2$ atmosphere incubator for 48 hours. At the end of the assay, 45 μl of freshly prepared XTT solution [0.5 mg/ml XTT sodium salt (2,3-Bis(2-methoxy-4-nitro-5-sulfophenyl)-2*H*-tetrazolium-5-carboxanilide inner salt, Sigma X4626) and 10 ng/ml PMS (1-Methoxy-5-methylphenazinium methyl sulfate, Sigma M8640) in culture media] were added to each well, and plates were cultured for 4 hours. Optical absorbances at 475 nm ($A_{475}$) and 660 nm ($A_{660}$) were measured spectrophotometrically at 20-22°C using a

Cytation 5 plate reader (Biotek). For each well, the specific absorbance $SA_{(test)}$ , the background-corrected specific absorbance $CSA_{(test)}$ , and the normalized specific absorbance $NSA_{(test)}$ were calculated according to the formulas:

(1) $SA_{(test)} = A_{475(test)} - \overline{A_{475(mock)}} - A_{660(test)}$ , where $\overline{A_{475(mock)}}$ is the average of all $A_{475}$ readings for the mock wells (column 12 of plate).

(2) $CSA_{(test)} = SA_{(test)} - \overline{SA_{(mock)}}$ , where $\overline{SA_{(mock)}}$ is the average of all *SA* for the mock wells (column 12 of plate).

(3) $NSA_{(test)} = CSA_{(test)} / \overline{CSA_{(no\ compound)}}$ , where $\overline{CSA_{(no\ compound)}}$ is the average of all *CSA* for the wells containing cells with no compound treatment (column 11 of the plate).

The drug effect and the drug concentration for 50% growth inhibition (GI50) were calculated using CompuSyn software (ComboSyn, Inc) (1, 2).

**Clonogenic cell survival assay.** Exponentially growing cell cultures were treated with compounds for 72 hours. At the end of treatment cell cultures were at 70-80% confluency. Cells were trypsinized and counted using trypan blue exclusion. 300 viable cells were re-plated in 100 mm cell culture plates in triplicate and allowed to form colonies for 14 days. At the end of the assay, culture media were removed, plates were washed twice with phosphate-buffered saline (PBS) and incubated for 10 min with 0.1% w/v crystal violet (Acros Organics 229641000) in 50% v/v methanol and 50% v/v phosphate-buffered saline (PBS, Invitrogen 21600044). Plates were excessively washed with tap water until colonies were clearly visible, and colonies were manually counted.

**Cell and tumor protein extraction.** For protein extraction all experimental procedures were performed on ice. Attached cells were washed once with cold PBS and lysed for 15 min with PTY buffer [50 mM HEPES, 50 mM NaCl, 5 mM EDTA, 1% Triton X-100, 50 mM NaF, 10 mM $Na_4P_2O_7$, 1 mM $Na_3O_4V$, 10 μg/ml phenylmethanesulfonyl fluoride, supplemented with protease and phosphatase inhibitors (Sigma P8340, P0044, and P5726)] (3). Cells were scrapped from culture plates with a cell lifter, passed through a 25-gauge hypodermic needle 10 times, and lysis was completed by incubation on ice for an additional 15 min. Cell debris was pelleted by centrifugation at 20,000 rcf for 20 min at 4°C. For tumor samples, 25 mg of tissue was homogenized in 500 μl T-PER buffer (Thermo Scientific 78510) supplemented with protease and phosphatase inhibitors. For homogenization, a THQ Digital Tissue Homogenizer (Omni International 12-500) and Stainless-Steel Generator Probe (Omni International B5-075) were used for a 30 sec burst at 20,000 rpm. Tissue homogenates were cleared by centrifugation at 10,000 rcf for 5 min. Total protein in lysates and homogenates was quantified using Pierce BCA (Thermo Scientific 23227) according to the manufacturer's instructions. All samples were stored at -80°C.

**Immunoblotting**. 10-20 μg total protein per well were resolved in NuPage Bis-Tris gels (Invitrogen), transferred on Immobilon-P PVDF membrane (Millipore), and stained with Ponceau S solution (Sigma P7170). Membranes were blocked with 5% w/v bovine serum albumin (BSA, Sigma A7284) or 5% w/v Blotto non-fat dry milk (NFDM, Santa Cruz Biotechnology SC-2324) dissolved in PBS, 0.05% v/v Tween20 (Sigma P7949). Conditions for western immunoblotting are given in **Supplementary Table 17**. Secondary antibodies were HRP-conjugated goat anti-rabbit or goat anti-mouse IgG (BioRad 1706516 and 1706515, respectively). Chemiluminescence was developed with SuperSignal West Pico (Thermo Scientific PI34078). The chemiluminescence signal was captured either digitally using a UVP Gel Imaging System (Analytik Jena US) or a Foto/Analyst FX 6-7206 (Fotodyne), or on autoradiography film (USA Scientific 1968-3810) which was developed in an SRX-101A Film Processor (Konica) and then scanned. For band intensity quantification, the ImageJ (version 1.52o, NIH) gel analysis tool was used on raw (unedited) images, the band intensity was quantified three times and the average was used in subsequent calculations.

**Preparation of compounds for *in vivo* use.** All compounding was performed aseptically. Solutions were filter-sterilized through a 0.2 μm pore filter (GE Healthcare 67802502 or Millipore-Sigma SE1M179M6).

For intraperitoneal administration of rapamycin, a stock solution was prepared by dissolving 7.5 mg rapamycin (Selleck Chemicals S1039) in 1 ml DMSO and stored aliquoted at -20°C. Injectable rapamycin solution (0.3 mg/ml) was freshly prepared by dissolving 40 μl of the 7.5 mg/ml stock solution in 960 μl sterile normal saline (Becton Dickinson) containing 0.25% v/v Tween-80 (Sigma P8074) and 0.25% w/v PEG300 (Sigma 202371) or in 960 μl filter-sterilized corn oil (Sigma C8267).

For oral administration of volasertib, a stock solution was prepared by dissolving 16 mg volasertib in 0.1 N HCl and stored aliquoted at -20°C. Ingestible volasertib solution (2.5 mg/ml) was freshly prepared by dissolving 156 μl of the 16 mg/ml stock volasertib solution in 844 μl 0.5% w/v hydroxyethyl cellulose (Natrosol 250).

For intraperitoneal co-administration of volasertib and rapamycin, the injectable solution (0.3 mg/ml rapamycin, 1.7 mg/ml volasertib) was freshly prepared by dissolving 40 μl of 7.5 mg/ml rapamycin stock and 1.7 mg volasertib in 960 μl corn oil.

**Short-term lung colonization studies.** Eight-week-old female CB17 SCID (B6.CB17-*Prkdc*$^{scid}$/SzJ, The Jackson Laboratory) were pre-treated for two days with rapamycin (1 mg/kg ip), volasertib (25 mg/kg po), or vehicle controls. 24 hours after the last dose, mice were anesthetized by isoflurane inhalation (5% v/v in $O_2$) and inoculated by tail-vein injection with $2x10^5$ ELT3-luciferase cells (ERL4) (4) in PBS at a total volume of 0.1 ml. Bioluminescence was measured 15 min after ip injection with XenoLight D-Luciferin (PerkinElmer 122799) using the Xenogen IVIS Spectrum In Vivo Imaging System (PerkinElmer). Bioluminescence image data were obtained at 1 h (baseline), 6 h and 24 h post cell inoculation. Animals that received poor luciferin administration, as determined during bioluminescence imaging, were excluded from analysis.

**Xenograft studies**. ELT3 and ELT3-245 cells were suspended in PBS at a final concentration of $2x10^7$ cells per ml. 621-101 cells were suspended in 50% v/v Matrigel (Corning 354234), 50% v/v PBS at a final concentration of $5x10^7$ cells per ml. Eight-week of female CB17 SCID mice (Taconic CB17SC-F EF) were inoculated with 0.1 ml cell suspension by subcutaneous injection in each of both flanks. Tumor width (W) and length (L) were measured using digital calipers, and tumor volume (V) was calculated using the formula [V = (W 2 × L) / 2] (5). For both intraperitoneal injections and oral gavage, the administration volume was 10 ml/kg of body weight. For rapamycin, mice were treated three times a week (Monday, Wednesday, and Friday). For oral volasertib, mice were treated twice a week on two consecutive days (Monday and Tuesday). For intraperitoneal co-administration of rapamycin and volasertib, mice were treated three times a week (Monday, Wednesday, and Friday). At endpoint, mice were euthanized by $CO_2$ narcosis followed by thoracotomy and removal of the heart and lungs.

**Gene expression studies.** RNA isolation and microarray gene expression assays were conducted by the UTHSC Molecular Resource Center (MRC) Institutional Core. Total RNA was isolated from tumor samples that were preserved in RNAlater Stabilizing Solution (Invitrogen-ThermoFisher). 350 μl RLT buffer (Qiagen) was added to 8-10 mg of sample and tissue disruption was performed in a TissueLyser (Qiagen) using 5 mm diameter stainless steel beads (Qiagen), and homogenates were cleared by centrifugation at 20,000 rcf and 4°C for 3 min. The cleared homogenate was used for RNA purification using the RNeasy Mini QIAcube Kit (Qiagen) in a QIACube instrument (Qiagen). RNA was quantified in a Synergy H1 multi-mode plate reader using a Take3 microvolume plate (Biotek). Samples with $OD_{260}/OD_{230} < 1.8$ were subjected to linear acrylamide / ethanol precipitation to increase purity. RNA integrity was assessed on a 2100 Bioanalyzer System using an RNA 6000 Nano Kit (Agilent). Microarray gene expression assays were performed from 1 μg total RNA on Clariom S rat-specific arrays (Affymetrix). Arrays were scanned on a GeneChip Scanner (Affymetrix).

Microarray gene expression data analysis was conducted by the UTHSC Molecular Bioinformatics (mBIO) Institutional Core. Fold-change (FC) for gene expression was calculated using standard methods. A FC threshold of 1.5 was applied to identify differentially expressed genes. A Welch *t* test and Benjamini – Hochberg false discover rate (FDR) were used to calculate statistical significance. Genes with an adjusted *P* value < 0.05 were considered statistically significant, and genes with an FDR < 0.05 were considered for further analysis. Pathway analysis was performed using Ingenuity Pathway Analysis (IPA) software (Qiagen).

For gene expression studies by RT-qPCR, the SuperScript VILO cDNA Synthesis Kit (Invitrogen) was used to synthesize cDNA from 1 μg total RNA. cDNA was used for target amplification using the TaqMan Fast Advanced Master Mix (Applied Biosystems) and gene specific TaqMan assays (Applied Biosystems, **Supplementary Table 18**) in technical triplicates. *Actb* and *Gapdh* were used as housekeeping gene. Amplification and detection were performed in an ABI 7500 Fast Real-Time PCR System. Pairwise fold-change

(FC) and fold-regulation were calculated using $RT^2$ Profiler (Qiagen). FC values <0.5 or >1.5 were considered significant. Fold-regulation (FR) was calculated as follows: for FC≥1, FR=FC, and for FC<1, FR=1/FC.
